# *Chlamydia muridarum* infection differentially changes smooth muscle contractility and responses to prostaglandins in uterus and cervix

**DOI:** 10.1101/448340

**Authors:** Jia Ming Lee, Jemma R. Mayall, Anne Chevalier, Dirk Van Helden, Jay C. Horvat, Philip M. Hansbro, Philip Jobling

**Affiliations:** School of Biomedical Sciences and Pharmacy, University of Newcastle, Callaghan, NSW, Australia; Hunter Medical Research Institute, New Lambton Heights, NSW, Australia; Priority Research Centre for Healthy Lungs, University of Newcastle, NSW, Newcastle Australia; Centenary Institute and the University of Technology Sydney, NSW, Australia

## Abstract

*Chlamydia trachomatis* infection is a primary cause of reproductive tract diseases including chronic pelvic pain and infertility. Previous studies showed that this infection alters physiological activities in mouse oviducts. Whether this occurs in the uterus and cervix has never been investigated. This study characterized the physiological activity of the uterus and the cervix in a *Chlamydia muridarum (Cmu)* mouse model of reproductive tract infection. Uterine or cervix smooth muscle contractility, responses to oxytocin or prostaglandins (PGF2α and PGE_2_) and mRNA expression of oxytocin and PG receptors were assessed 14 days post infection. *Cmu* infection did not affect the contractions of the uterine horn but significantly decreased the contraction amplitude of the cervix. *Cmu* infection did not alter the responses of uterine horn or cervix to oxytocin, however PGF2α induced contractions of the uterine horn, but not the cervix, were significantly increased following *Cmu* infection. PGE_2_ contraction amplitude in both the uterine horn and cervix was unaffected by *Cmu* infection. An upregulation of *Ptgfr* and a down-regulation of *Ptegr4* mRNA expression was observed in the uterine horn following *Cmu* infection. These results indicate that *Cmu* infection alters contractility and prostaglandin signalling in the female reproductive tract but the effects are localised to specific regions.

## Author summary

*Chlamydia trachomatis* infection is one of the most commonly reported sexually transmitted infections (STIs) worldwide. The infection is readily treated with antibiotics; however, majority of the patients do not display symptoms meaning that they often miss early treatment for acute infection. This enables the spread of bacteria to the upper reproductive tract and the development of diseases including chronic pelvic pain, pelvic inflammatory disease and subsequent infertility. Recent studies reported altered physiological contraction in the oviducts of *Chlamydia* infected mice. This directly affects the transportation of oocyte in the oviduct. It is not known if this phenomenon extends to the lower part of the reproductive tract such as the uterus and cervix which has implications for sperm transport and ascension of pathogens towards the oviduct. Here we demonstrate that *Chlamydia* infection of mice significantly reduced the spontaneous contractile activity in the cervix and altered the response of the uterus to some endogenous signalling molecules (prostaglandins). These results show for the first time that chlamydia infection can alter smooth muscle activity along the reproductive tract in a region-specific manner.

## Introduction

Between 2012 to 2014, *Chlamydia trachomatis* infection was the most frequently reported sexually transmitted infection in Australia [1]. Whilst in England and the United States, *Chlamydia trachomatis* infection is among the most prevalent sexually transmitted infection reported [2, 3]. It is a major concern worldwide as it not only leads to clinically important morbidities, it also causes significant financial burden to individuals and healthcare services [4]. *Chlamydia* infections can be easily cured in the early stages with antibiotics. However, untreated infected patients are either asymptomatic or often misinterpreted the abdominal discomfort as dysmenorrhea (menstrual pain) or general abdominal pain [5]. Subsequently, ascension of the bacteria to the upper reproductive tract occurs and it often progresses into serious complications. Pelvic inflammatory disease is one of the common complications following *Chlamydia* infection. It is a constellation of upper genital tract inflammations that causes serious long-term effects in patients, including tubal factor infertility, ectopic pregnancy [6, 7] and chronic pelvic pain [8, 9] that affect quality of life [10].

Recently it was shown that *Chlamydia* infection disrupts the pace-maker activity of mouse oviducts, thus, affecting the spontaneous contraction of these tissues. This disrupted motility is a potential contributor to tubal infertility. Whether this extends to the lower region of the female reproductive tract (FRT) remains unknown. The uterus and cervix, just like the oviducts, contract spontaneously in the non-pregnant state in human and mouse [11-14]. Spontaneous contraction in the FRT is regulated by period of the menstrual cycle and is believed to have an essential role in maintaining fertility of women. Although the exact function of FRT contraction has not been fully elucidated, what is already known is that it helps in oocyte propulsion and menstrual shedding [15], sperm transportation [16] and perhaps is also involved in the transportation or shedding of pathogens. Apart from spontaneous contraction, smooth muscles along the FRT are also responsive to endocrine, paracrine and neuronally released signalling molecules. The hormone oxytocin increases uterine [17-20], cervical and vaginal motility [14]. Prostaglandins signalling is recognized in modulating uterine motility [21]. Furthermore, the importance of prostaglandins and mast cell mediators released by degranulating cells [22] induced by *Chlamydia* infection in other tissues [23] is also being recognised. Importantly, these mediators also induce the contraction of the FRT [24] and are suggested to be associated with pain in dysmenorrhea [25].

In other viscera, notably the bladder and bowel, inflammation or infection are known to alter smooth muscle function. This occurs in inflammatory bowel disease (IBS) [26, 27], ulcerative colitis [28], interstitial cystitis [29], urinary tract infection [30] and cystic fibrosis [31]. Inflammatory mediators exert their effects by directly acting on smooth muscle cells or, indirectly, through stimulating the release of mediators from other cells such as mast cells. Although there has not been investigations of whether immune response affect the normal physiology and contractility along the FRT, evidence from preterm and term deliveries suggest that immune mediators regulate myometrial contraction during parturition. According to studies, inflammatory neutrophils and macrophages are observed in the uterus, decidua and cervix in both clinical and pre-clinical models during labour process. These immune cells, coupling with mast cells and chemokines, coordinate the timely contraction of the uterus, cervical ripening and dilation and rupture of the fetal membranes during parturition. Furthermore, these cells have also been linked to the occurrence of pre-term labour [32-35]. *Chlamydia* infection alters the immune profile of the FRT [36-39] but the effects on motility are unknown.

We hypothesized that the immune changes triggered by *Chlamydia* infection might alter the responses of the FRT to various endogenous mediators and affect its motility along the FRT. To replicate *Chlamydia* reproductive tract infection in human, we infected mice intravaginally with *Cmu* [40]. This is a mouse-adapted strain that induces upper reproductive tract inflammation characterized by the development of hydrosalpinx similar to human *C. trachomatis* reproductive tract infection [36, 37, 39, 40]. We investigated the effects of infection on the spontaneous contraction of mouse uterine horn and cervix. Responses of these tissues to endogenous mediators such as oxytocin and prostaglandins were assessed.

## Results

### Pathology of *Cmu* infection

*Cmu* infection caused a significantly higher cross-sectional area in the oviducts of mice, signifying inflammation and swelling. The cross sectional areas of the left oviduct of sham-inoculated and *Cmu*-inoculated mice were 1.169±0.15 mm^2^ and 3.081±0.45 mm^2^, respectively (*n*=8, *P*=0.0013, unpaired *t*-test); and of the right oviduct were 0.993±0.19 mm^2^ and 3.159±0.42 mm^2^ (*n*=8, *P*=0.0004, unpaired *t*-test, S1 Fig). To confirm productive *Cmu* infection in the upper reproductive tract, we quantified *Cmu* ribosomal 16S RNA in the left uterine horn. There was no detectable expression in the sham-inoculated uterine horn while a detectable threshold level of 16S was observed in the uterine horn of 14 days post-inoculation (dpi) in *Cmu*-inoculated mice (relative expression 1.562±0.75, S2 Fig).

### Effects of *Cmu* infection on spontaneous contraction

To investigate whether *Cmu* infection affects contraction in the uterine horn and cervix, we first compared the spontaneous contraction (baseline contraction) of both the uterine horn and cervix between sham-inoculated and *Cmu*-inoculated mice. *Cmu* infection did not alter spontaneous contraction in terms of amplitude and frequency in the uterine horn (*n*=13-14) (Fig 1A, B). The amplitude and frequency of baseline contraction in the uterine horn were 47.02±5.00% and 8.08±0.65 per 5 min for sham-inoculated compared with and 44.84±2.86% and 7.50±0.44 per 5 min for *Cmu*-inoculated mice. *Cmu* infection also did not alter the responses of the uterine horn to 60 mM KCl which made this suitable as an internal control (Fig 1C). In contrast, the amplitude of baseline contraction of the cervix was significantly decreased in infected mice (*n*=14; P =0.0245, unpaired *t*-test, Fig 1D), being 36.51±6.57% and 18.36±3.83% for sham-inoculated and *Cmu*-inoculated mice, respectively. The contractile frequency of the cervix was also decreased by infection, being 6.79±1.86 per 5min in sham-inoculated and 2.86±0.95 per 5min in *Cmu*-inoculated mice, though this was not quite reach statistical significance (*n*=14, P =0.0546, Mann Whitney U test, Fig 1E). Infection did not alter cervix responses to 60 mM KCl (Fig 1F).

**Fig 1:**
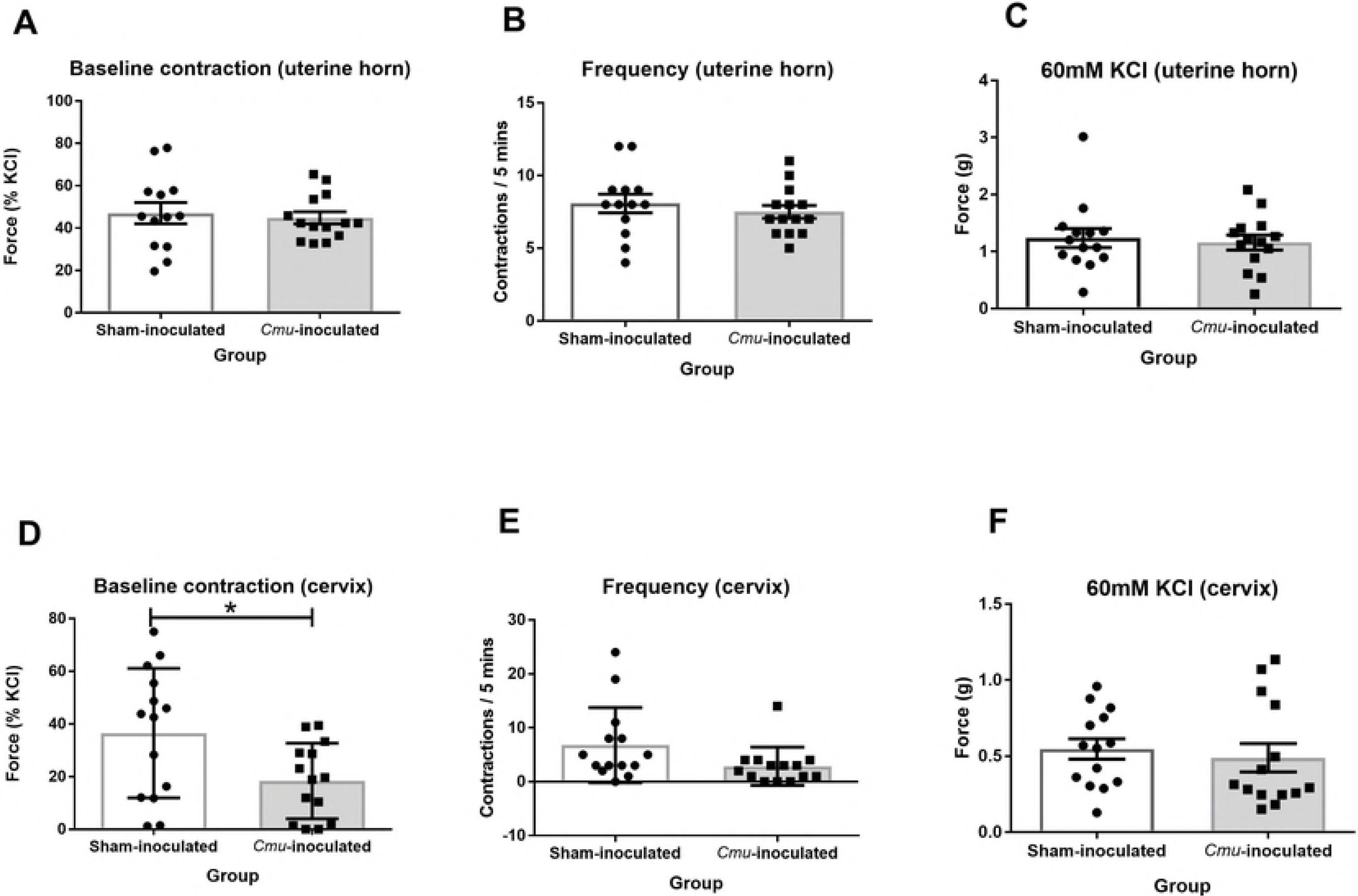
The amplitude and frequency of baseline contraction in the uterine horn and cervix of sham-inoculated and *Cmu*-inoculated mice. (A-C) 14 dpi *Cmu* infection did not affect the spontaneous contraction of the uterine horn and its response to 60 mM KCl. (D) *Cmu* infection significantly reduced the force of baseline contraction in the cervix (\**P*<0.05, unpaired *t*-test); (E-F) while did not alter its baseline frequency and the response to 60 mM KCl. Data are presented as mean ± S.E.M. (*n* = 14).

### Effects of *Cmu* infection on mediator-induced contractions

#### The responses to oxytocin

We next investigated the responses of the uterine horn and cervix to oxytocin (S3 Fig). The maximum response (E_max_) of contractile amplitude and frequency to oxytocin was not statistically different in the uterine horn of sham-inoculated versus *Cmu*-inoculated mice (Fig. 2A, B). There was also no statistical difference in the EC_50_ of oxytocin in the uterine horns between the two groups. The EC_50_ of oxytocin for sham-inoculated was 21.70±4.75 nM and 43.59±17.71 nM for *Cmu*-inoculated mice (*P*=0.5887; Mann-Whitney test). The cervix of *Cmu*-inoculated mice showed heightened contraction amplitude to increasing dosage of oxytocin, but it was not significantly different from the sham-inoculated. The EC_50_ of oxytocin in both groups was not significantly different either (120.20±39.71 nM in sham-inoculated, 102.50±35.25 nM in *Cmu*-inoculated mice; *P*=0.7463, unpaired *t*-test). Similarly, the contraction frequency in the cervix to oxytocin was not affected by *Cmu* infection too.

**Fig 2:**
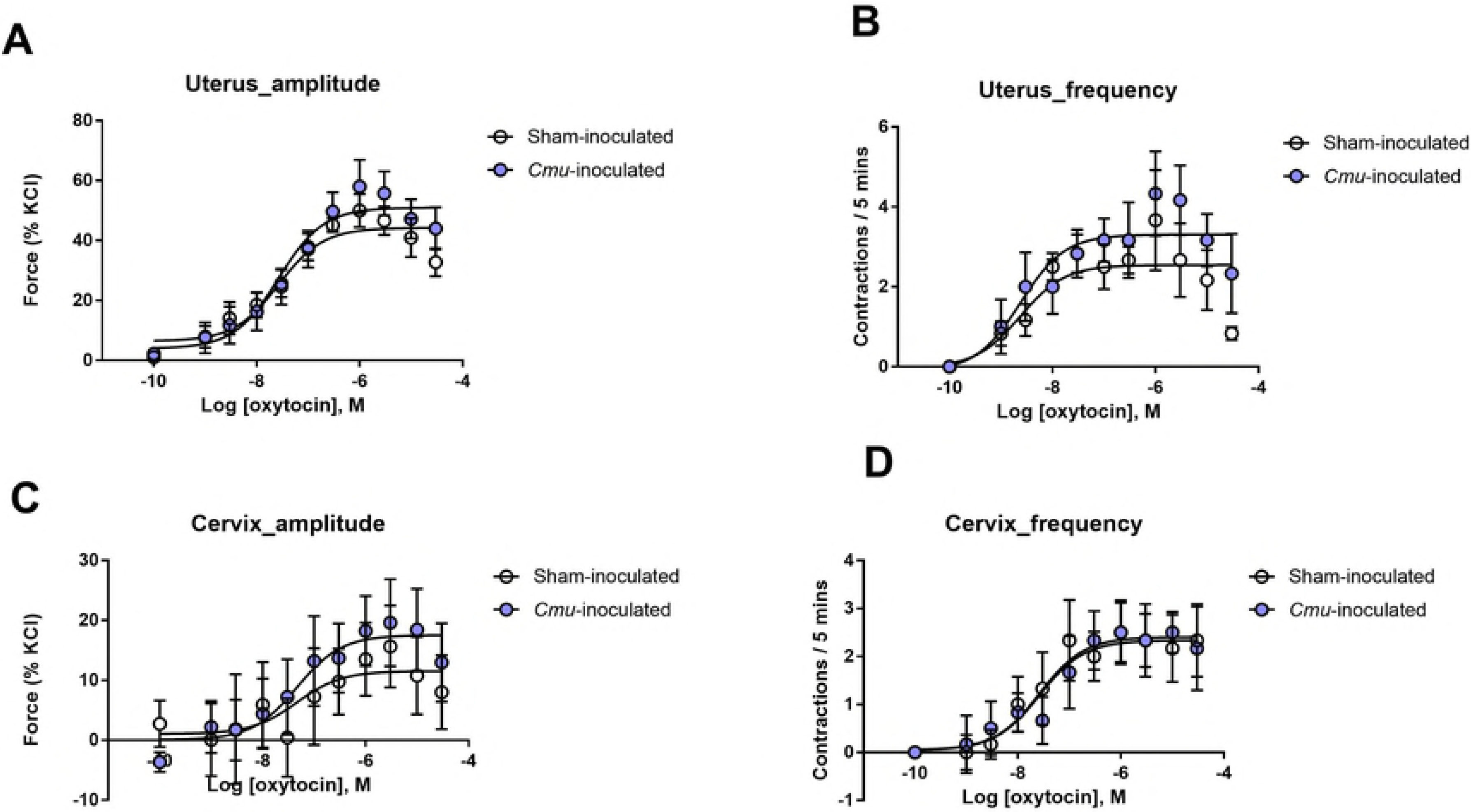
The comparison of responses of the uterine horn and cervix to oxytocin in sham-inoculated and *Cmu*-inoculated mice. (A-B) Higher E_max_ in the contractile amplitude and frequency was detected in the uterine horn of *Cmu*-inoculated mice, but it was not significantly different between the two groups. (C) The E_max_ of contractile amplitude to oxytocin was nonsignificantly increased in the cervix of *Cmu*-inoculated mice. (D) Contractile frequency of the cervix was not affected by *Cmu*-infection. Data are presented as mean ± S.E.M (*n*=6; unpaired *t*-test or Mann-Whitney test).

#### The responses to PGs

The contractile responses of the uterine horn and cervix to PGF2α and PGE_2_ in *Cmu* infection were assessed (S4 Fig). Addition of PGF2α potentiated spontaneous contractions in the uterine horn in both sham-inoculated and *Cmu*-inoculated mice (Fig 3A). Uterine horns of *Cmu*-inoculated mice had significantly higher responses to 1 μM PGF2α compared to sham-inoculated controls. The contractile amplitude at this concentration after corrected for baseline contractions was 15.59±7.23% and 50.62±9.72% for sham-inoculated and *Cmu*-inoculated mice, respectively (*n*=7-8; *P*=0.0108; One-way ANOVA with Dunnett as post-hoc, Fig 3B). Similarly, the exposure to 3 μM PGF2α elicited heightened contraction in the uterine horn of *Cmu*-inoculated mice too, and the difference of the contractile amplitude after baseline correction was significant between the sham-inoculated and *Cmu*-inoculated (sham-inoculated – 4.52±5.22%; *Cmu*-inoculated mice – 26.19±6.34%; *P*=0.0195; One-way ANOVA with Dunnett as post-hoc; Fig 3B).

**Fig 3:**
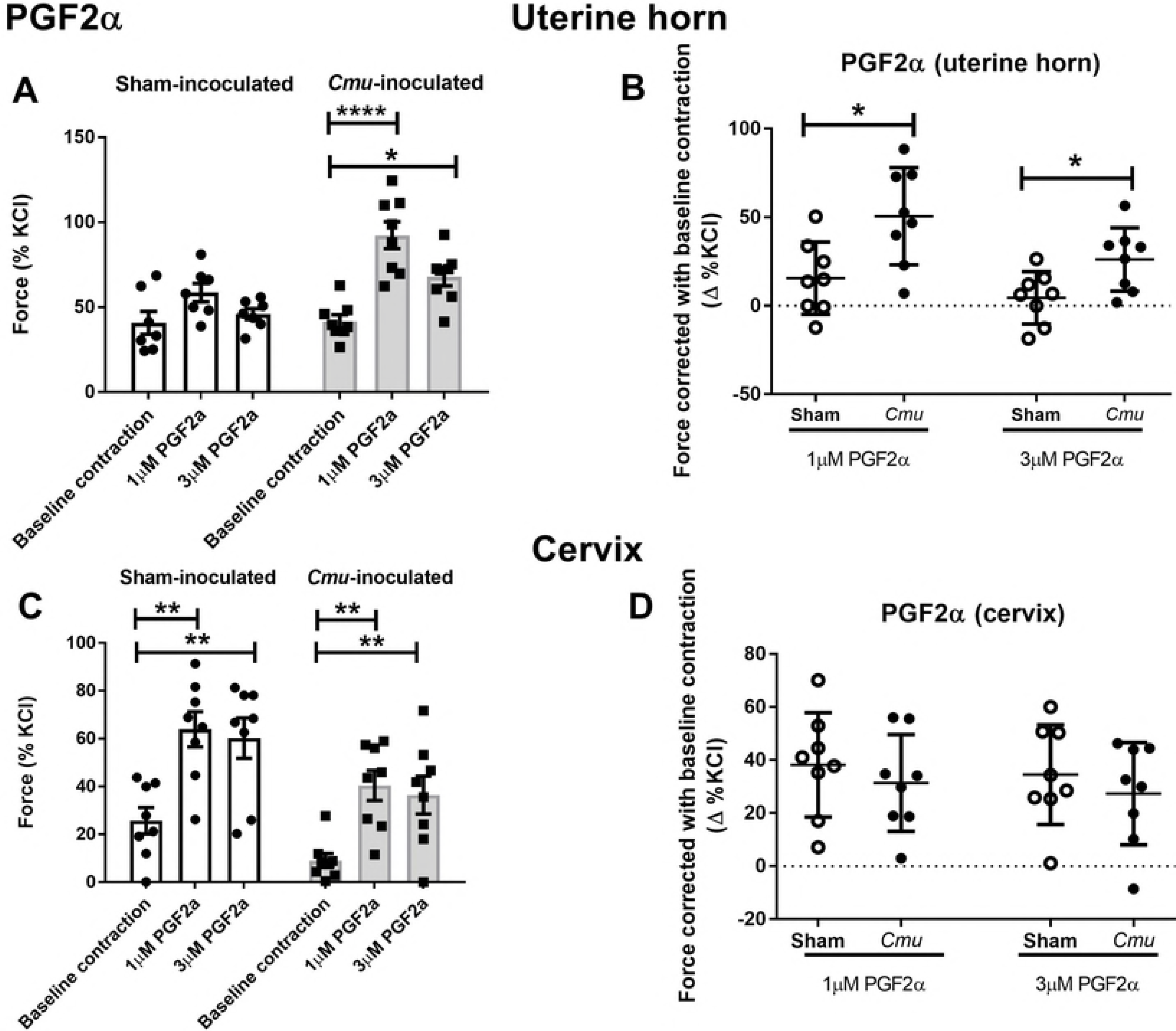
PGF2α significantly increased contraction force in the uterine horn, but not in the cervix of *Cmu*-inoculated mice. PGF2α increased contractile amplitude of the uterine horn (A) and cervix (C) of the progesterone-primed mouse. 14 dpi *Cmu* infection significantly increased contractile amplitude in the mouse uterine horn (B), but not in the cervix (D). Data are presented in mean ± S.E.M (*n* = 8; one-way ANOVA with Dunnett’s post-hoc; \**P*<0.05; \*\**P*<0.01; \*\*\*\**P*<0.0001).

PGF2α also stimulated contractions in the cervix of both sham-inoculated and *Cmu*-inoculated mice (Fig 3C). However, after baseline correction, there was no significant difference between the two groups (at 1μM – sham-inoculated = 38.24±6.95%, *Cmu*-inoculated mice = 31.37±6.46%, *P*=0.8873; at 3μM – sham-inoculated = 34.53±6.64%, *Cmu*-inoculated mice = 27.36±6.82%, *P*=0.8741, Fig 3D).

There was no significant differences in the baseline-corrected contractile amplitude to PGE_2_ in both the uterine horn (*P*=0.7949; Fig 4B) and cervix (*P*=0.5933; Fig 4D) of both groups. Nevertheless, we observed opposing trend elicited by PGE_2_ in both the uterine horn and cervix. PGE_2_ decreased contractile amplitude in the uterine horn (Fig 4B) while increased that in the cervix (Fig 4D). This has never been characterized elsewhere and the opposing responses to PGE_2_ in these two entities could have some roles in maintaining the homeostasis of FRT that is yet to be discovered.

**Fig 4:**
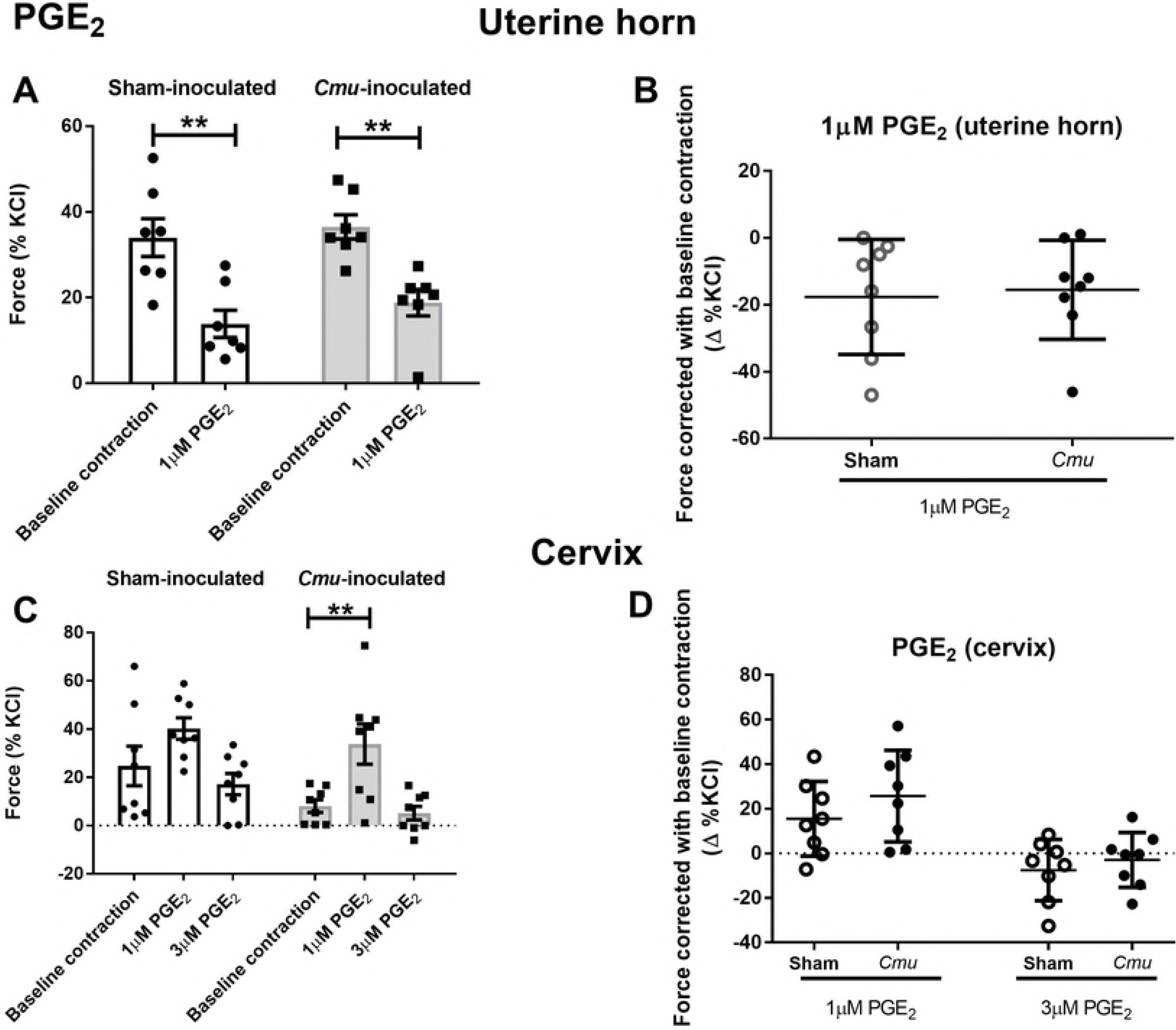
The contractile amplitude of mouse uterine horn and cervix to PGE_2_ in sham-inoculated and *Cmu* infection. PGE_2_ exhibited opposing effects in the uterine horn and cervix of progesterone-primed mouse. Contractile amplitude to 1μM PGE_2_ was significantly reduced in the uterine horn (A) but increased in the cervix (C). The responses of both the uterine horn and cervix to PGE_2_ was not affected by 14 dpi *Cmu* infection (B, D). Data are presented in mean ± S.E.M (*n* = 8; one-way ANOVA with Dunnett’s post-hoc; \**P*<0.05; \*\**P*<0.01).

#### qPCR expression of receptor mRNA

In order to compare pharmacological responses to potential changes in receptor expression we examined receptor mRNA in the uterine horns using qPCR. The receptor genes for selected agonists are: OTR: oxytocin receptor; *Hrh-1:* histamine H_1_ receptor; *Ptgfr:* PGF2α receptor; *Ptger1 – 4:* receptors for PGE_2_, namely EP_1_ EP_2_, EP_3_ and EP_4_. There were no significant differences in the mRNA expression of *OTR*, *Hrh-1*, *Ptger1*, *Ptger2* or *Ptger3* between sham-inoculated and *Cmu*-inoculated mice (Fig 5A-B, D-F). A significantly higher expression of *Ptgfr* was observed in the uterine horn of *Cmu*-inoculated compared to sham-inoculated mice (relative expression of sham-inoculated – 0.0254±0.0.0019, *Cmu*-inoculated – 0.0430±0.0.0051, *n*=12; P =0.0039, Fig 5C). In contrast, *Ptegr4* was downregulated in the uterine horn of *Cmu*-inoculated compared to sham-inoculated mice (relative expression of sham-inoculated – 0.4167±0.0277; *Cmu*-inoculated mice – 0.2935±0.0260, *n*=12; P =0.0038, Fig 5G).

**Fig 5:**
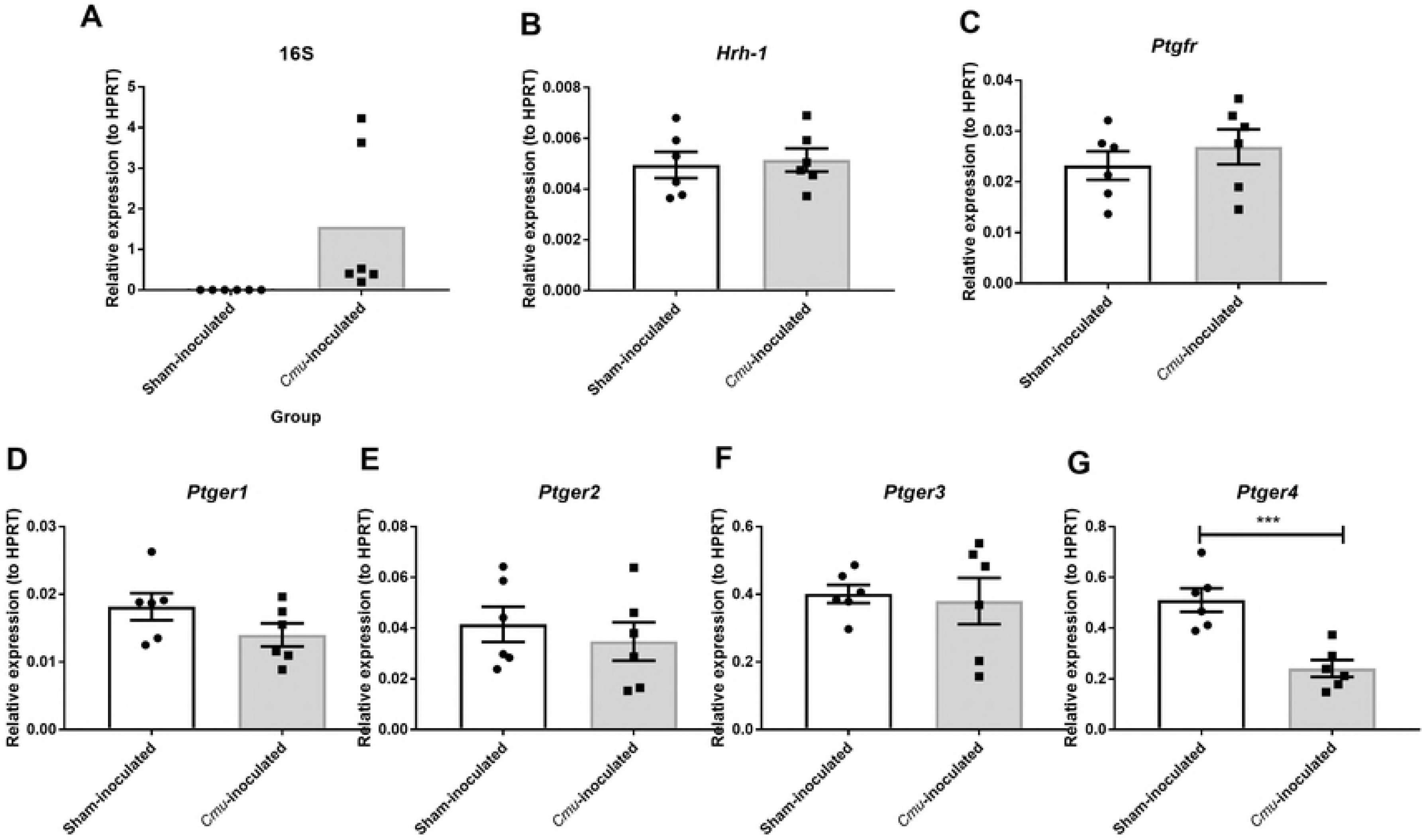
qPCR quantification of receptor mRNA expression in the uterine horn of sham-inoculated and *Cmu*-inoculated mice. *Cmu* infection significantly upregulated downregulated the mRNA expression of *Ptgfr* and *Ptger4* in mouse uterine horn, respectively (G) (\*\**P*<0.01). *OTR* was also downregulated in the uterine horn of *Cmu*-inoculated mice but it was not significant (A). \*\**P*<0.0001; NS-non-significant. (Abbreviations: *OTR*-oxytocin receptor gene; *Hrh-1*-histamine receptor H1 gene; *Ptgfr* – prostaglandin F receptor gene; *Ptgerl* – prostaglandin E receptor 1 gene; *Ptger2* – prostaglandin E receptor 2 gene; *Pter3* – prostaglandin E receptor 3 gene; *Ptger 4* – prostaglandin E receptor 4 gene). Data are presented in mean ± S.E.M (*n* = 12; unpaired *t*-test).

## Discussion

It has long been recognized that *Chlamydia* reproductive tract infections in women elicit a range of immunological responses in infected areas [41]. Despite the alarming increase in *Chlamydia* reproductive tract infections, little is understood if such immunological changes affect the physiological functions of the female reproductive tract. Based on studies conducted on mice, *Chlamydia* infection causes loss of pace-making ability in the oviduct that eventually impedes the transportation of the oocyte to the uterus, resulting in oviduct occlusion [42, 43]. However, to date, there have been no studies that have investigated whether sexually transmitted infection-associated pathology extends to the uterus or the lower reproductive tract.

Here, we focused on the uterus and cervix, as these organs are the main routes for microbe ascension that lead to subsequent complications [16].

We discovered that a 14 day course of *Cmu* infection did not alter contractile amplitude and frequency of spontaneous contraction of the uterine horn, signifying the pace-making ability of the uterine horn was still intact. The physiological reason for ongoing uterine motility in the non-pregnant state is to assist with sperm transport, or acts to maintain the uterine environment in a state consistent with implantation. A single, short-term *Cmu* infection usually does not pose a risk to long term complications like pelvic inflammatory disease and infertility in women [44], which perhaps explains the maintained motility in the uterine horn in our model. More perplexing was our finding that *Cmu* infection significantly reduced the contractile amplitude of spontaneous contractions in the cervix, but the contractile frequency was not affected. The function of cervical contraction in non-pregnant women is poorly understood [45]. Some studies suggest that it facilitates sperm transport. It has also been suggested that cervical contractions are partly responsible for the spread of infection to the upper region [16, 46], in which case it can be speculated that a reduced contractile amplitude would be protective. *Cmu* infection changes the microbiome composition in the FRT [47], such change could modify the spontaneous contractility of smooth muscle [48, 49]. Recent studies suggests that the microbiome in the body could regulate smooth muscle contractility. This is proven in mouse intestinal smooth muscle where a change in the guts microbiota inhibits the contractile amplitude of spontaneous phasic contractions in colonic longitudinal muscle, while not affecting the contractile frequency [48]. This is parallel with our finding in the cervix where *Cmu* infection reduced the contractile amplitude of spontaneous contraction while keeping the contractile frequency intact. Further studies are needed to explore the mechanism behind and its impact on pathologies such as pre-term labour and miscarriage.

Although spontaneous contractions were largely unaffected, many endogenous mediators contribute to uterine motility *in vivo.* We first investigated the response of the uterine horn and cervix to oxytocin, the prominent stimulant for female reproductive tract motility. The analysis of dose-response relationships did not show statistical differences in the uterine horn and cervix between *Cmu*-inoculated and sham-inoculated mice. Two indices, E_max_ and EC_50_, however, displayed trends towards higher values in *Cmu* infection. When we analysed *OTR* mRNA expression we found no differences, although there were trends towards reduced expression. There are several interleukin (IL) response elements flanking both the rat and human OTR promoter region, of which nuclear factor (NF)-IL6 response elements is one of them [50, 51]. Although we found no consistent evidence for IL-6-induced changes in OTR expression, previous studies have implicated inflammation and IL-6 in particular changes *OTR* expression. A study by Ma and co-workers showed that IL-6 elevated uterine *OTR* mRNA expression and binding capacity of the receptor [52]. While another study showed that IL-6 increased *OTR* mRNA and protein expression in pregnant uterine tissue but had no effect on non-pregnant uterine tissue in rat [32]. Interestingly, a study conducted by Schmid and co-workers on myometrial cells display negative regulation of *OTR* gene expression by IL-1β and IL-6 treatment [50]. Clearly the link between inflammation and changes to *OTR* signalling is complex.

We did observe increases in baseline contractions of the uterine horn in response to increasing doses of oxytocin regardless of the treatment condition. This is in line with previous findings where increased baseline contraction to increasing oxytocin concentration was reported in the uterus of pregnant and non-pregnant animal models [53, 54]. This phenomenon, however, was absent in the cervix in our study. Cervical strips exhibited phasic contraction throughout the oxytocin experiment. There have been no investigations yet on the dose-response relationship of oxytocin on non-pregnant longitudinal cervical strips. Hence, this is the first report demonstrating the differences between the physiological response of uterine horn and cervix in mice.

It is established that increased pro-inflammatory mediators such as IL-1, IL-6, IL-8 and tumour necrosis factor alpha (TNFα) have direct or indirect effects on the contractility of the female reproductive tract. This has been reported in pre-term and term pregnancies where these mediators play vital roles in stimulating necessary processes for parturition, including myometrial contraction and cervical ripening [35]. Up-regulation of genes that encode IL-lβ, Il-6, IL-8 and TNFα were observed in patients with dysmenorrhea, and increased production of these cytokines may induce multiple actions that lead to primary dysmenorrhea [52]. Furthermore, these immune mediators also promote the cyclooxygenase pathway [55] to induce prostaglandin release, as well as degranulating mast cells that lead to the release of uterotonic mediators [22]. In *Chlamydia* reproductive tract infection, increases in IL-6 and IL-8 are reported [41]. Whether they stimulate uterine hypercontractility leading to dysmenorrhea remains unknown. This underpins the second part of our study, which is to investigate if there are changes in the contractile intensity in the uterine horns and cervix in response to endogenous mediators.

The responses of uterine horns and the cervix in *Cmu* infection to prostaglandins were explored. There was a marked increase in contractile amplitude to 1 μM PGF2α in the uterine horns of *Cmu*-inoculated mice compared to sham-inoculated controls, but this was not observed in the cervix. For PGE_2_, no significant difference between the sham-inoculated and *Cmu*-inoculated mice was observed in either tissue. Prostaglandins mediate their effects via FP and EPs receptors. Specifically, PGF2α has the highest binding affinity to the FP receptor encoded by *Ptgfr.* It also has an affinity for EP receptors encoded by *Ptger1* to 4. PGE_2_ acts mainly *via* the four subtypes of EP receptors, namely EP_1_ EP_2_, EP_3_ and EP_4_ receptors encoded by *Ptger1-4*[56, 57]. Our findings demonstrated that the expression of *Ptgfr* in the uterine horns was increased by *Cmu* infection. In addition, *Ptger4* that encodes EP_4_ receptor was significantly downregulated in *Cmu* infection. The activation of FP receptor mediates myometrium contraction while the activation of EP_4_ receptor causes smooth muscle relaxation [58]. Upregulation of *Ptgfr* and the downregulation of *Ptger4* in the uterine horns is consistent with our functional result where *Cmu* infection increased the amplitude of uterine horn contractions to PGF2α. Increased responses to PGF2α in the uterine horn has long been known to contribute to primary dysmenorrhea [59]. It is uncertain in our models if such increases in *Cmu* infection generates discomfort or symptoms of abdominal pain. Also, we do not know if the increased response of the uterine horn to PGF2α would disrupt the normal physiological functions of the reproductive tract in *Cmu* infection.

The activation of EP_4_ receptors generally promotes anti-inflammatory processes in the viscera. It reduces chronic inflammation *in vivo* in inflammatory bowel syndrome, reduces duodenal and intestinal inflammation as well as allograft rejection [60, 61], and suppresses the release of pro-inflammatory cytokines in experimental model of myocardial ischemia, allograft rejection in cardiac transplantation, colitis and abdominal aortic aneurysm [60-62]. In the female reproductive tract, the exact association of EP_4_ receptor with inflammatory processes is not understood. Given the observations in our study, the reduced *Ptger4* could be a factor that causes prolonged inflammation in the upper reproductive tract in *Cmu* infection. Further investigation is required to delineate the pathogenesis of upper reproductive tract inflammation in *Cmu* infection.

Apart from the findings focusing on *Cmu* infection, we detected an interesting observation of PGE_2_ on the uterine horn and cervix. PGE_2_ exerted opposing effects on these tissues in progesterone-primed mice, regardless of the infection status. The effects of PGE_2_ are dependent on which of the four receptor subtypes are stimulated in a given situation. It triggers contraction *via* EP_1_ and EP_3_ receptors while inducing relaxation *via* EP_2_ and EP_4_ receptors [58].

The mice in our study were pre-treated with progesterone prior to group allocation. The results suggest that progesterone treatment up-regulates EP_2_ and EP_4_ in the mouse uterine horn and up-regulates EP_1_ and EP_3_ in the cervix since both the organs display the opposite contraction trend in response to PGE_2_. This is corroborated by a previous study, where progesterone treatment increases EP_2_ gene expression in the mouse uterus [63]. Nonetheless, ours is novel evidence as previous studies focused only on the molecular expression of EP and FP receptors and there is lack of functional characterisation of the physiological effects. Furthermore, it is also the first evidence to show the differential response in the mouse uterine horn and cervix in response to PGE_2_.

In summary, we demonstrated that *Cmu* infection did not affect the spontaneous contraction of uterine horn in mice, but it markedly decreased that in the cervix. Responses to oxytocin were minimally altered in the uterine horn in *Cmu* infection while there was no changes observed in the cervix. We also showed a heightened response to PGF2α in the uterine horn in *Cmu* infection. This was correlated with the upregulation of *Ptgfr* and the downregulation of *Ptger4* mRNA expression that encodes the FP and EP_4_ receptor individually in the uterine horn of *Cmu*-inoculated mice. Furthermore the response to PGE_2_ differs along the reproductive tract. These novel functional findings also show the opposing effects of PGE_2_ in progesterone-primed uterine horn and cervix.

## Materials and methods

### Ethics Statement

All animal procedures were conducted in accordance with the Australian Code For The Care And Use Of Animals For Scientific Purposes 8^th^ Edition (2013) endorsed by the National Health and Medical Research Council (NHMRC), the Australian Research Council, the Commonwealth Scientific Industrial Research Organisation and Universities Australia. It was approved by The University of Newcastle Animal Care and Ethics Committee (A-2011-109).

### Mouse treatments

Female wild-type C57BL/6 mice between 18-21 weeks old were housed under specific pathogen-free conditions. At day 1, animals were subcutaneously injected with medroxprogesterone acetate (Pfizer, Australia; 2.5 mg in 200 μL saline) under isofluorane anaesthesia to synchronize them in the diestrus stage of the estrous cycle [36, 37, 39]. At day 8, mice were infected by intravaginal inoculation of either 5 × 10^4^ inclusion forming units of *Cmu* in 10 μL sterile sucrose phosphate glutamate for *Cmu*-inoculated or 10 μL of sucrose phosphate glutamate for sham-inoculated under ketamine-xylazine anaesthesia. This improves infection susceptibility under ketamine-xylazine anaesthesia [36, 37, 39]. Mice were monitored daily for 14 dpi. This period was sufficient for the infection to induce adaptive immune responses around the FRT that resembles chronic *Chlamydia* infection and the inception of hydrosalpinx in infected mice (unpublished data). Then, mice were sacrificed by sodium pentobarbitone overdose, and vaginal lavages were collected for estrous cycle confirmation as previously described [14, 39]. Left uterine horn was dissected out, trimmed of visceral fat, snap-frozen in liquid nitrogen (N_2_) and stored at −80°C until needed. Right uterine horn and cervix were placed in chilled physiological saline solution (PSS, 120 mM NaCl, 5 mM KCl, 2.5 mM CaCl_2_, 2 mM MgCl_2_, 25 mM NaHCO_3_, 1 mM NaH_2_PO_4_ and 1 mM glucose; gas with 95% O_2_ and 5% CO_2_).

### *Cmu* infection

The diameter of both oviducts were examined using a caliper, and the cross sectional area was calculated to determine hydrosalpinx. Swelling of the oviducts indicated inflammation resulting from upper reproductive tract infection.

The expression of *Cmu* ribosomal 16S RNA in the left uterine horn was quantified using quantitative polymerase chain reaction (qPCR) as previously described [64-68]. 16S ribosomal RNA indicates active infection and successful spread of *Cmu* to the upper reproductive tract [37, 69].

### RNA isolation, reverse-transcription PCR (RT-PCR) and qPCR

Frozen left uterine horns were thawed and homogenized in 500 μL of TRIzol^®^ (Invitrogen, Mount Waverly, VIC, Australia) using a Tissue-Tearor stick homogenizer (BioSpec Products, Bartesville, OK) on ice. RNA was extracted according to the manufacturer’s instructions (TRIzol^®^, Invitrogen, Mount Waverly, VIC, Australia) [36, 37, 39]. The end-product was reverse-transcribed to cDNA using M-MLV Reverse Transcriptase kit (Promega, Australia) and a T100™ Thermal Cycler (BioRad). qPCR was performed on a Mastercycler^®^ *ep* Realplex (Eppendorf, Germany) using SYBR reagents. mRNA expression was calculated using 2^−△△Ct^ relative to the reference gene hypoxanthine-guanine phosphoribosyl-transferase (HPRT) and expressed as relative expression. Primers used were as follows:

**Table 1.**
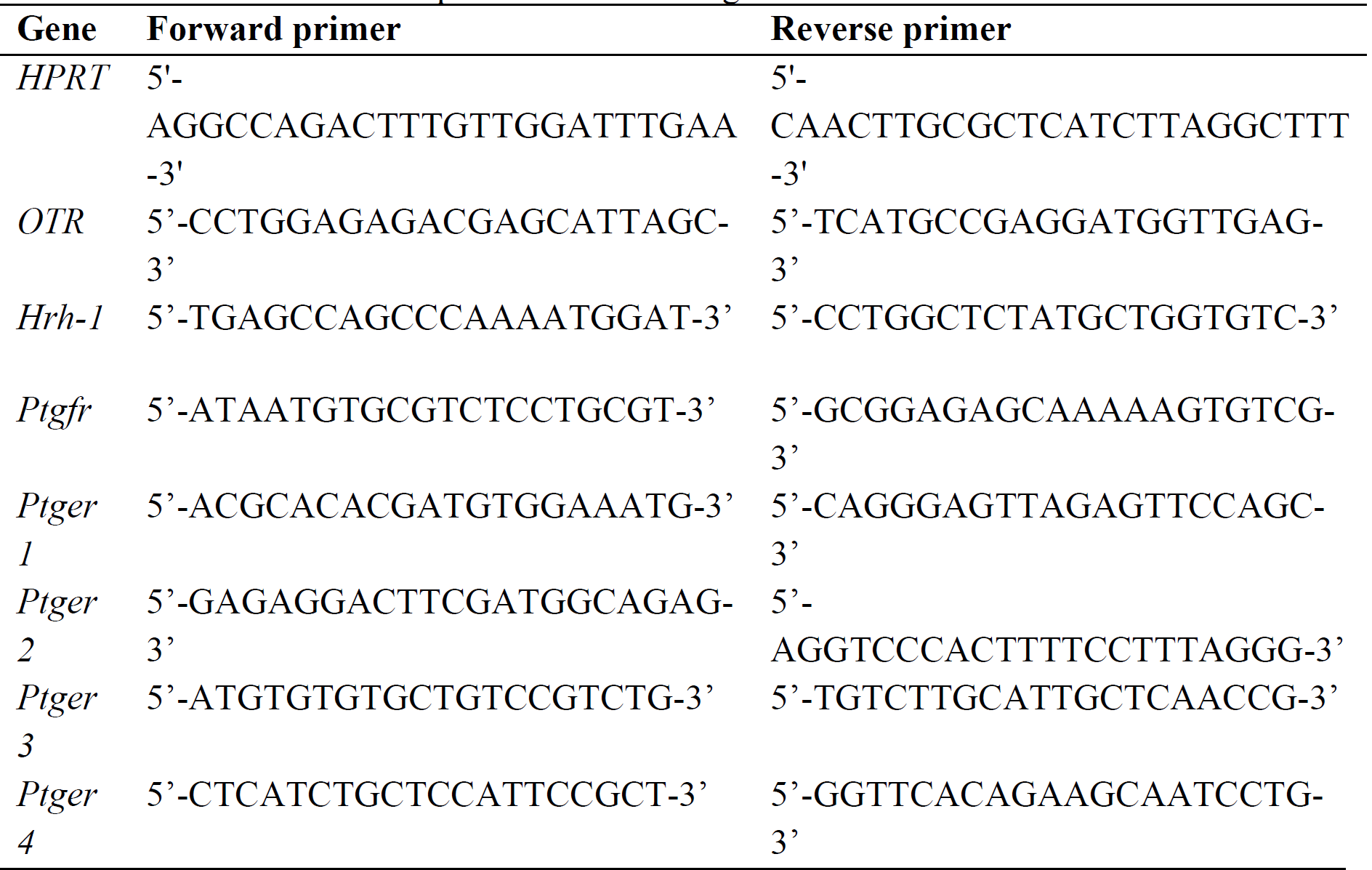
Forward and reverse primers of selected genes

### Preparation of myometrial and cervical strips

Components of the freshly dissected mouse female reproductive tract were isolated. Right uterine horns were trimmed of connective tissue and opened along the mesometrial border. A uterine strip of 1 cm in length measured from the end of oviduct was prepared. Cervixes were dissected out, trimmed of connective tissue, and opened circumferentially to expose the smooth muscle. The longitudinal uteri and cervical strips were attached at one end to a metal hook and the other end to a tension transducer (Grass FT03) in 4 mL organ baths. Tissues were then equilibrated under 10 mN (cervix) and 5 mN (uteri) for 30 min in PSS at 37°C before being challenged with submaximal concentrations of 60 mM KCl. Baseline contractions (spontaneous contraction without stimulation) and mediator-induced contractions were recorded as previously described [14]. At the end of the experiment, tissues were blotted dry with towel paper and wet weight measured.

### Contractile activity

Experiments involved 30 min equilibration, and 30 min baseline and drug-induced contractions. Oxytocin, PGF2α and PGE_2_ (Cayman Chemical, Ann Arbor, USA) were dissolved in dimethyl sulfoxide (DMSO). The final concentration of DMSO in PSS did not exceed 1:1000. Cumulative oxytocin dose responses were constructed by adding increasing concentrations into the organ bath every 5 min. In separate experiments, single concentrations of PGF2α (1 μM) and PGE_2_ (3 μM) were tested. The concentration for each mediator was selected based on previous studies [70, 71]. Each mediator was applied for 5 min before washing. Tissues were washed 3-5 times before the next drug was added. At the end of the experiment, tissues were contracted with 60 mM KCl again to ensure tissue viability. The contraction amplitude before and after the addition of respective mediators was measured. It was then normalized to baseline contraction and wet weight and was expressed as % of the 60 mM KCl response. Contractile frequency was measured as the amount of contractions that occurred in 5 min and expressed as contractions/5 min. Analysis was performed using LabChart 8 Reader (ADInstruments, Australia).

### Chemicals

Oxytocin, PGF2α and PGE_2_ (Cayman Chemical, Ann Arbor, USA) were dissolved in dimethyl sulphoxide (DMSO). The final concentration of DMSO in the PSS did not exceed 1:1000.

### Statistical Analysis

Normality test was performed using Statistical Package for the Social Science v24 software (SPSS Inc., Chicago, IL, USA) and data are presented as means ± standard error of the mean (S.E.M.). Statistical significance was determined using Graphpad Prism Software v6 (San Diego, CA) with significance set at P<0.05. Unpaired *t*-test was used for the comparison in S1, Fig 1B, 1C, 1D, 2 (for E_max_ and EC_50_), 4A, 4B, 4D and 5 as these data were normally distributed and fit the assumptions for parametric test. Non-parametric Mann-Whitney U test was used for data that were not normally distributed (Fig 1A, 1E, 1F). One-way ANOVA was used with Dunnett’s multiple comparison as *post-hoc* test for the comparison for PGF2_α_ (Fig 3) and the comparison for PGE_2_ (Fig 4C).

## Acknowledgements

We thank Mr. Peter Dosen for technical training and support for contractility assays.

## Supporting Information

**S1 Fig.**
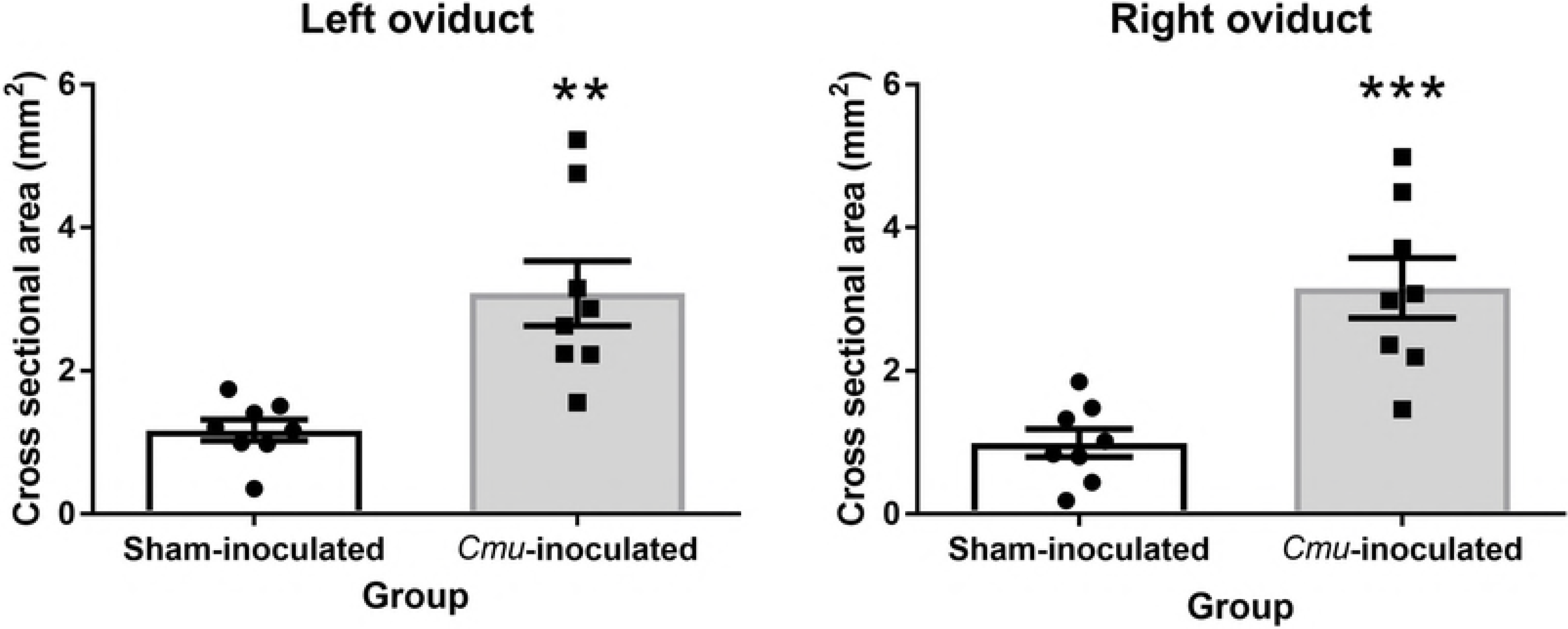
Intravaginal *Cmu* treatment increases cross sectional area of mouse oviducts. Higher cross sectional area (mm^2^) was observed in both the oviducts of mouse treated with *Cmu* (14 dpi) when compared to sham-inoculated. Data are expressed as mean ± S.E.M. (*n* = 8 per group) \*\**P*<0.01; \*\*\**P*<0.001, unpaired student *t*-test.

**S2 Fig.**
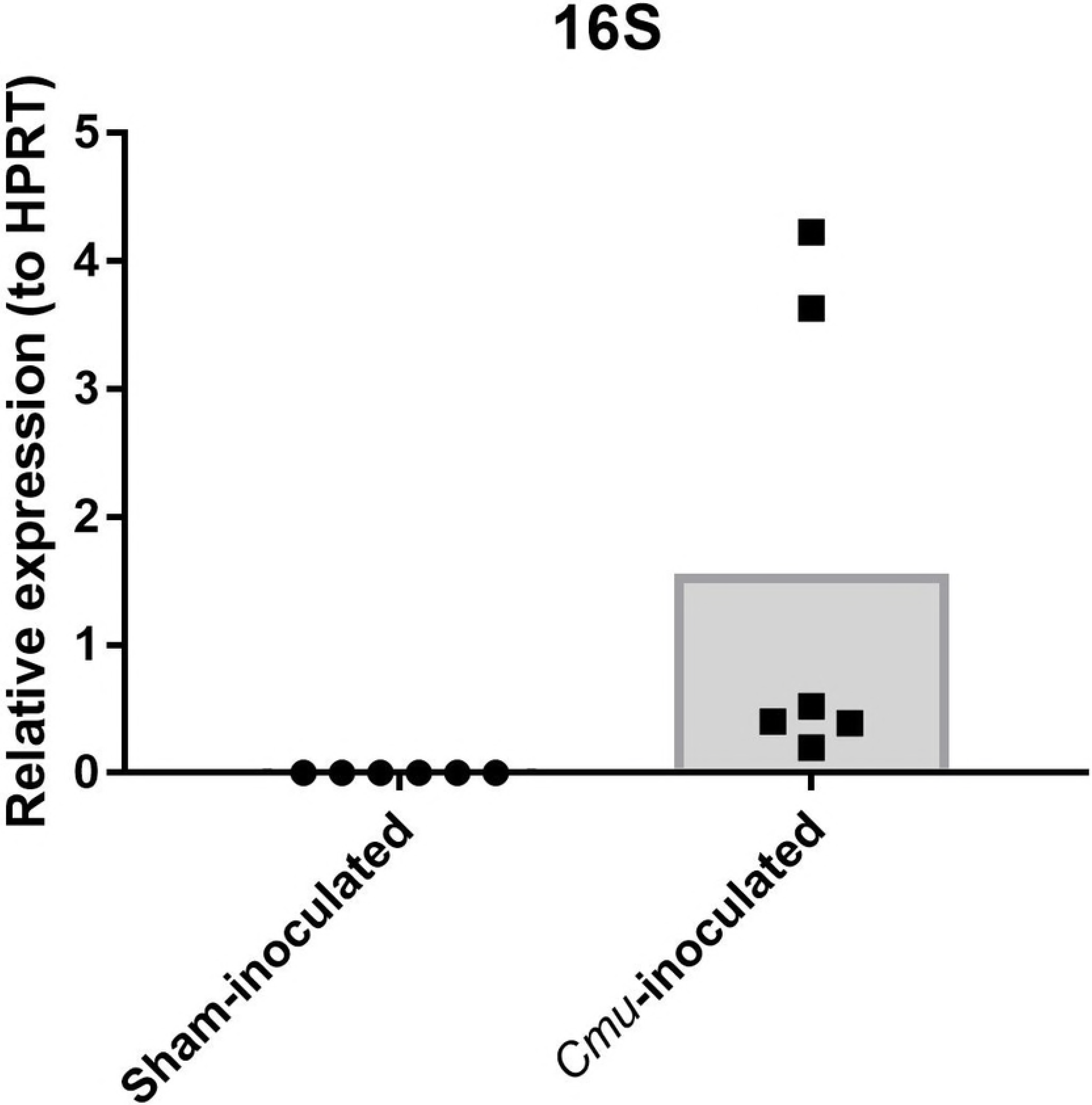
mRNA expression of 16S in the left uterine horn of sham-inoculated and *Cmu*-inoculated mice after 14 dpi. 16S expression was only detected in the left uterine horn of *Cmu*-inoculated mice and not in the sham-inoculated.

**S3 Fig.**
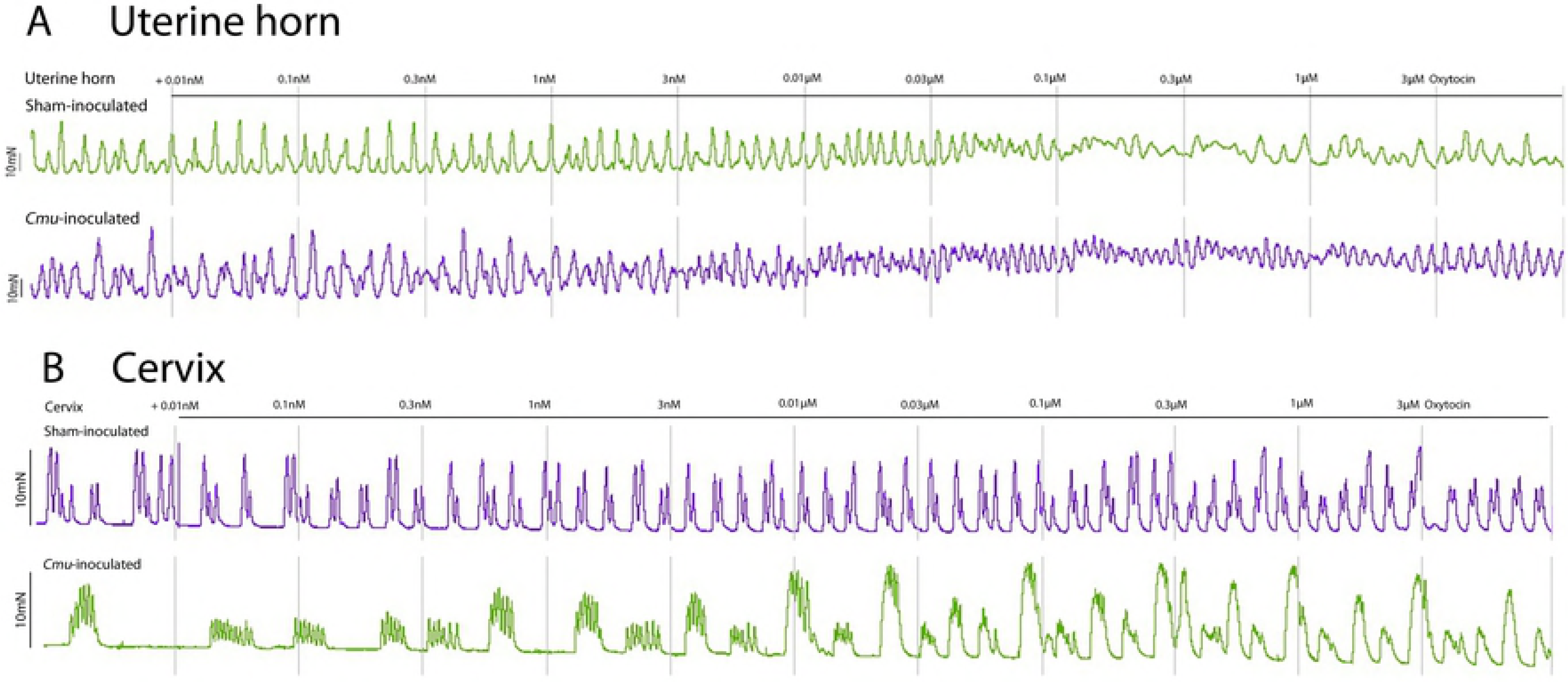
Contractile responses of the uterine horn (A) and cervix (B) to cumulative concentration oxytocin (0.01 nM to 3 μM) in sham-inoculated and *Cmu*-inoculated. Increased baseline contraction was observed in the uterine horn to increased oxytocin concentration, however, this was not observed in the cervix.

**S4 Fig.**
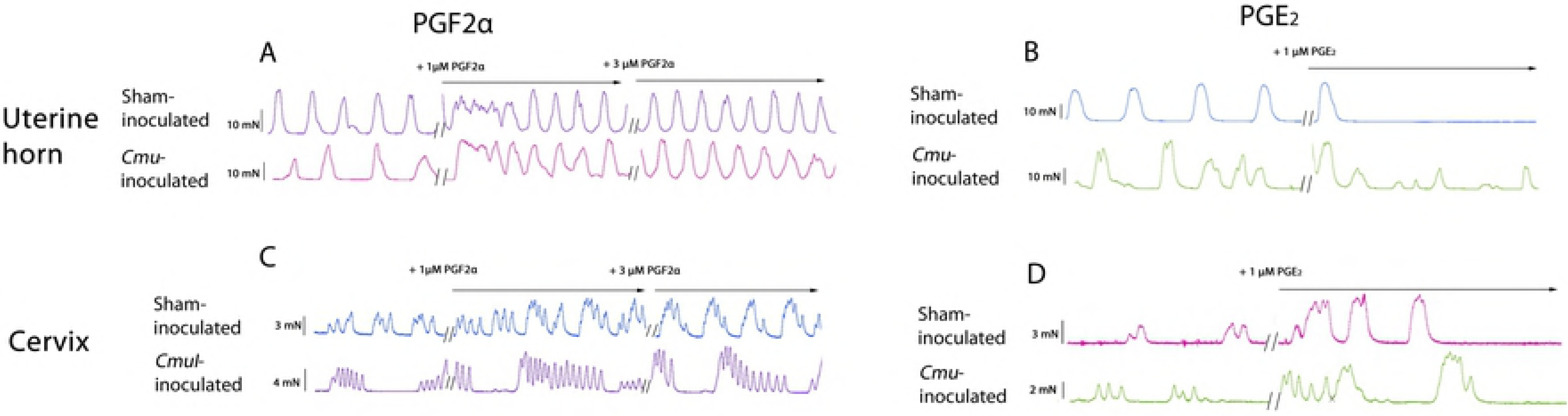
Contractile responses of the uterine horn and cervix to PGF2α and PGE_2_ in sham-inoculated and *Cmu*-inoculated. Addition of 1 μM PGF2α induced tonic contraction in the uterine horn (A) and higher contractile amplitude and frequency in the cervix of both sham-inoculated and *Cmu*-inoculated mice (C). PGE_2_ caused opposing effects in the uterine horn and cervix, inhibiting contraction in the uterine horn (B) while enhancing contractions in the cervix (D).

